# Quantitative proteomics of the CDK9 interactome reveals a novel function of the HSP90-CDC37-P-TEFb complex for BETi-induced HIV-1 latency reactivation

**DOI:** 10.1101/2022.01.20.477160

**Authors:** Cong Wang, Zhenrui Pan, Yaohui He, Zhanming Zhang, Rongdiao Liu, Yuhua Xue, Qiang Zhou, Xiang Gao

**Affiliations:** State Key Laboratory of Cellular Stress Biology and Fujian Provincial Key Laboratory of Innovative Drug Target Research, School of Pharmaceutical Sciences, Xiamen University, Xiamen, Fujian 361102, China; Department of Molecular and Cell Biology, University of California, Berkeley, CA 94720, USA

**Keywords:** HSP90, P-TEFb, HIV Latency Reactivation, Brd4, Quantitative Proteomics

## Abstract

Brd4 has been intensively investigated as a promising drug target because of its implicated functions in oncogenesis, inflammation and HIV-1 transcription. The formation of the Brd4-P-TEFb (CDK9/Cyclin T1) complex and its regulation of transcriptional elongation is critical for HIV latency reactivation and expression of many oncogenes. To further investigate the mechanism of the Brd4-P-TEFb complex in controlling elongation, mass spectrometry-based quantitative proteomics of the CDK9 interactome was performed. Upon treatment with the selective BET bromodomain inhibitor (BETi) JQ1, 535 proteins were successfully identified with high confidence as CDK9-interacting proteins. Among them, that increased bindings of HSP90 and CDC37 to CDK9 were particularly striking, and our data suggest that the HSP90-CDC37-P-TEFb complex is involved in controlling P-TEFb’s dynamic equilibrium during BETi-induced HIV-1 latency reactivation. Furthermore, the HSP90-CDC37-P-TEFb complex directly regulates HIV-1 transcription and relies on the recruitment by heat shock factor 1 (HSF1) for binding to the HIV-1 promoter. These results advance the understanding of HSP90-CDC37-P-TEFb in HIV-1 latency reversal and enlighten the development of potential strategies to eradicate HIV-1 using a combination of targeted drugs.

## Introduction

As a transcriptional co-activator and epigenetic reader, bromodomain-containing protein 4 (Brd4) binds to acetylated lysine residues of histone H3 and H4,^1-3^ and recruits positive transcription elongation factor b (P-TEFb), composed of cyclin-dependent kinase 9 (CDK9) and Cyclin T1, to phosphorylate the C-terminal domain (CTD) of RNA polymerase (Pol) II on Ser-2 and switch Pol II from promoter-proximal pause to transcriptional elongation.^4, 5^ This is critical for the reversal of HIV latency and expression of many oncogenes such as *c-myc, bcl-xl* and *pax5*, and as such makes Brd4 a promising therapeutic target. ^6, 7^

Most of the developed BET inhibitors (BETi) block the activity of Brd4 by competitively binding to the bromodomains of Brd4 and disrupting its interactions with acetylated lysine.^8^ Furthermore, the inhibitors can also be designed as hetero-bifunctional PROTAC (Proteolysis Targeting Chimera) probes to recruit the E3 ubiquitin ligase for selective degradation of Brd4 and inhibition of its functions.^9, 10^ In addition, several BETi including JQ1 have been shown to effectively reactivate latent HIV through antagonizing Brd4 to promote the formation of the Tat-Super Elongation Complex (Tat-SEC).^11, 12^

CDK9 is vital for HIV-1 and cellular gene transcription.^13, 14^ The dynamic equilibrium of P-TEFb is critical for BETi-induced HIV latency reversal. Besides the Tat-SEC complex, to determine whether P-TEFb is also recruited by other proteins to gene promoters when Brd4 is inhibited, we quantitatively identified the CDK9 interactome under the JQ1 treatment conditions by using the stable-isotope labelling by amino acids (SILAC) method that was integrated with liquid chromatography mass spectrometry (MS).^15, 16^ Through this quantitative approach and subsequent biological validation, we found that more HSP90-CDC37-P-TEFb complex could be formed upon the treatment with JQ1.

Heat shock protein (HSP90) is one of the most abundant proteins in human cells and plays important roles in stabilization and maturation of client proteins including key kinases involved in cell cycle control and oncogenic transformation.^17^ Our earlier research demonstrates that HSP90 and its partner CDC37 are required for proper folding/stabilization of CDK9 in the production of mature P-TEFb.^18^ Recently, the role of HSP90 in HIV-1 latency reactivation has also been revealed. It was shown that HSP90 could be localized to the viral promoter ^19, 20^ and participated in the NF-κB-dependent transactivation pathway.^21^ Here, we demonstrate that upon the inhibition of Brd4, the formation of the HSP90-CDC37-P-TEFb complex increased significantly and is subsequently recruited by heat shock factor 1 (HSF1) to the viral LTR to reactivate latent HIV-1 transcription. Our results indicate this complex as directly involved in controlling the dynamic equilibrium of P-TEFb.

## Results

### Quantitative identification of CDK9 interactome upon JQ1 treatment using SILAC

A mass spectrometry (MS)-based quantitative proteomics approach involving the stable-isotope labeling by amino acids (SILAC) in cell culture was used to determine the CDK9-associated proteins after Brd4 was inhibited by JQ1 (Figure 1A). Briefly, F1C2 cells, in which CDK9-Flag is expressed to about the same level as endogenous CDK9,^22^ were cultured in a specific medium containing “heavy” SILAC amino acids for at least seven passage until cells were completely incorporated with heavy isotopic amino acids (^13^C_6_)-labeled Arginine and Lysine). The isotopically labeled cells and the control F1C2 cells grown in normal media were then treated with JQ1 and DMSO, respectively. CDK9-Flag and its associated proteins were immunoprecipitated from nuclear extracts of an equally mixed sample of “heavy” and normal F1C2 cells and then digested and analyzed by liquid chromatography-tandem mass spectrometry (LC-MS/MS). Using this procedure, we identified 535 proteins with high confidence and reproducibility from both independent biological replicates (Figure 1B, Figure S1A, Extended Data-1, Supporting Information).

**Figure 1.**
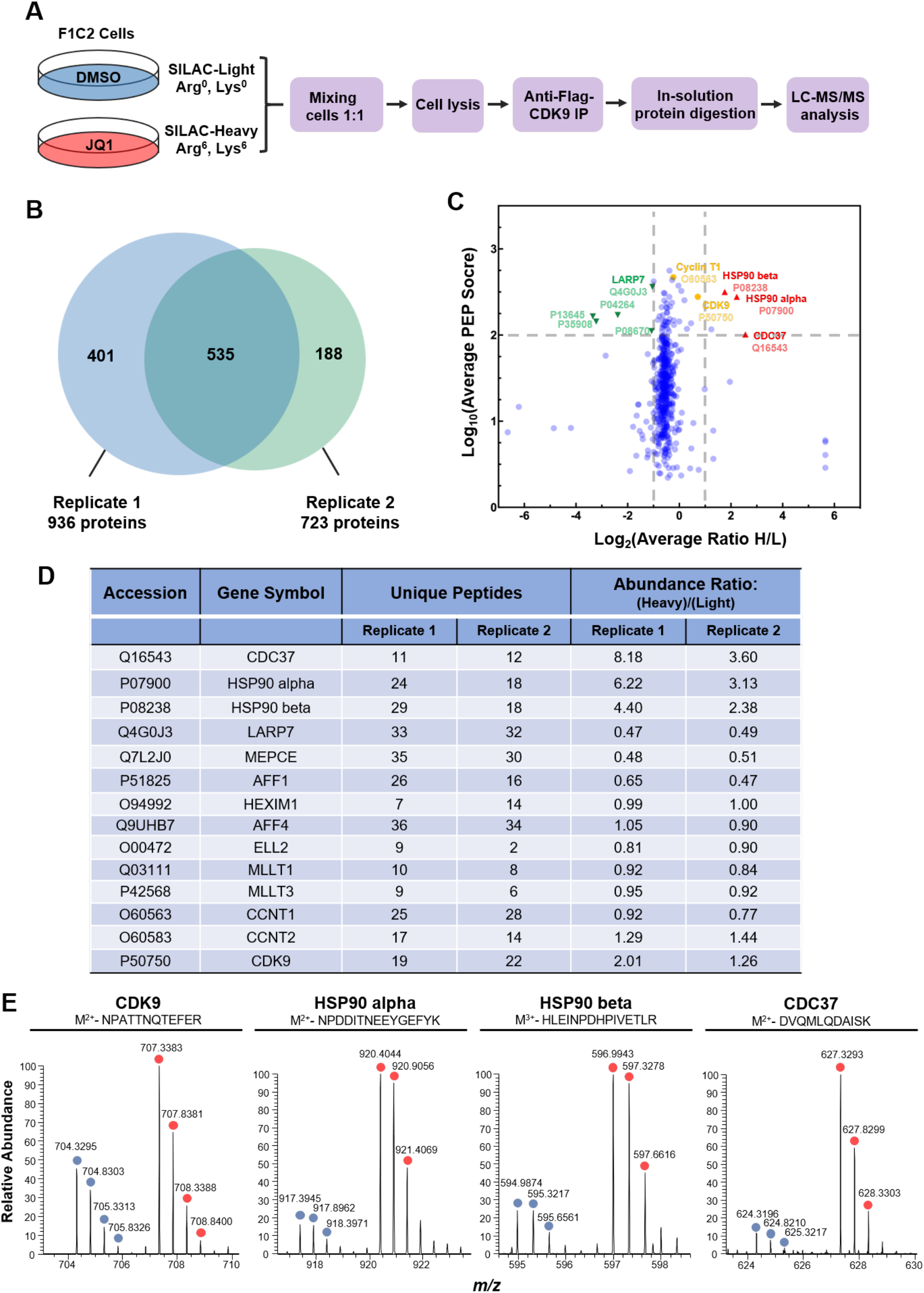
Identification of CDK9 interactome under treatment of BET inhibitor using SILAC proteomics. **A**. Schematic diagram of SILAC proteomics for quantification of CDK9-associated proteins. For protein quantification, F1C2 cells cultured in media containing stable isotopic amino acids Arg^6^ and Lys^6^ were treated with 1 μM JQ1 for 12 h. CDK9-associated proteins were analyzed by liquid chromatography-tandem mass spectrometry (LC-MS/MS). **B**. The Venn diagram showing the overlap of CDK9-associated proteins quantified in two biological replicates. **C**. Volcano plot illustrated the log_2_ SILAC ratios determined in two biological replicates and their PEP score (log_10_) calculated by Proteome Discover software (version 2.4). CDK9 and Cyclin T1 were labeled as yellow. Proteins with increased binding to CDK9 were labeled as red, and proteins with decreased binding to CDK9 were labeled as green. **D**. Quantitative results of the representative CDK9 interacting proteins (ratios= Heavy (JQ1) /Light (DMSO)). **E**. Mass spectra of the unique peptides for quantification of CDK9 (aa 346-358), HSP90 alpha (aa 490-499), HSP90 beta (aa 331-337), and CDC37 (aa 111-121), respectively.

Next, the identified proteins were subjected to ontology enrichment analysis. The top 20 enriched molecular functions were prominently associated with gene transcription, translation, RNA splicing/binding, mRNA metabolic process, protein folding and ATPase activity (Figure S1B, Supporting Information), which were consistent with previous research.^23^ Figure 1D summarizes the quantitative results of key CDK-interacting proteins for controlling gene transcription. Notably, LARP7 and MePCE, two components of the 7SK snRNP, showed significantly decreased interactions with CDK9 upon JQ1 treatment (Figure 1C&D),^24^ and this result is consistent with our previous study.^11^ In contrast, the components of the Super Elongation Complex (SEC) including AFF4, ELL2 and ENL, displayed no obvious changes (Figure 1D).

### CDK9 displays increased interactions with HSP90 and CDC37 upon treatment with JQ1

Remarkably, HSP90-alpha, HSP90-beta and CDC37 showed a significant increase in bindings to CDK9 upon exposure to JQ1 (Figure 1C&D). The quantitative ratios were manually confirmed using both the full-mass spectra and MS/MS tandem mass spectra of representative unique peptides of CDK9, HSP90-alpha, HSP90-beta and CDC37 as shown in Figure 1E (Extended Data-2, Figure S1C, D, E&F, Supporting Information). The increased interactions of HSP90 and CDC37 with CDK9 as revealed by our quantitative proteomic analyses suggest that the HSP90-CDC37-P-TEFb complex probably plays a critical role in controlling cellular gene transcription when the Brd4 function is blocked.

To further verify the interaction between CDK9 and the HSP90-CDC37 complex, we performed anti-Flag immunoprecipitation (IP) in F1C2 cells, which was followed by Western blotting (WB). Consistent with the above SILAC proteomic analysis MS analyses, the CDK9-Flag-associated HSP90 and CDC37 were dramatically increased following JQ1 treatment (Figure 2A) and in a dose- and time-dependent manner (Figure 2B&C). Furthermore, the treatment with a different BET inhibitor iBET-151 also increased the association of HSP90 and CDC37 with CDK9 (Figure 2D). Taken together, these data confirmed the increased formation of the HSP90-CDC37-P-TEFb complex under the BETi treatment conditions.

**Figure 2.**
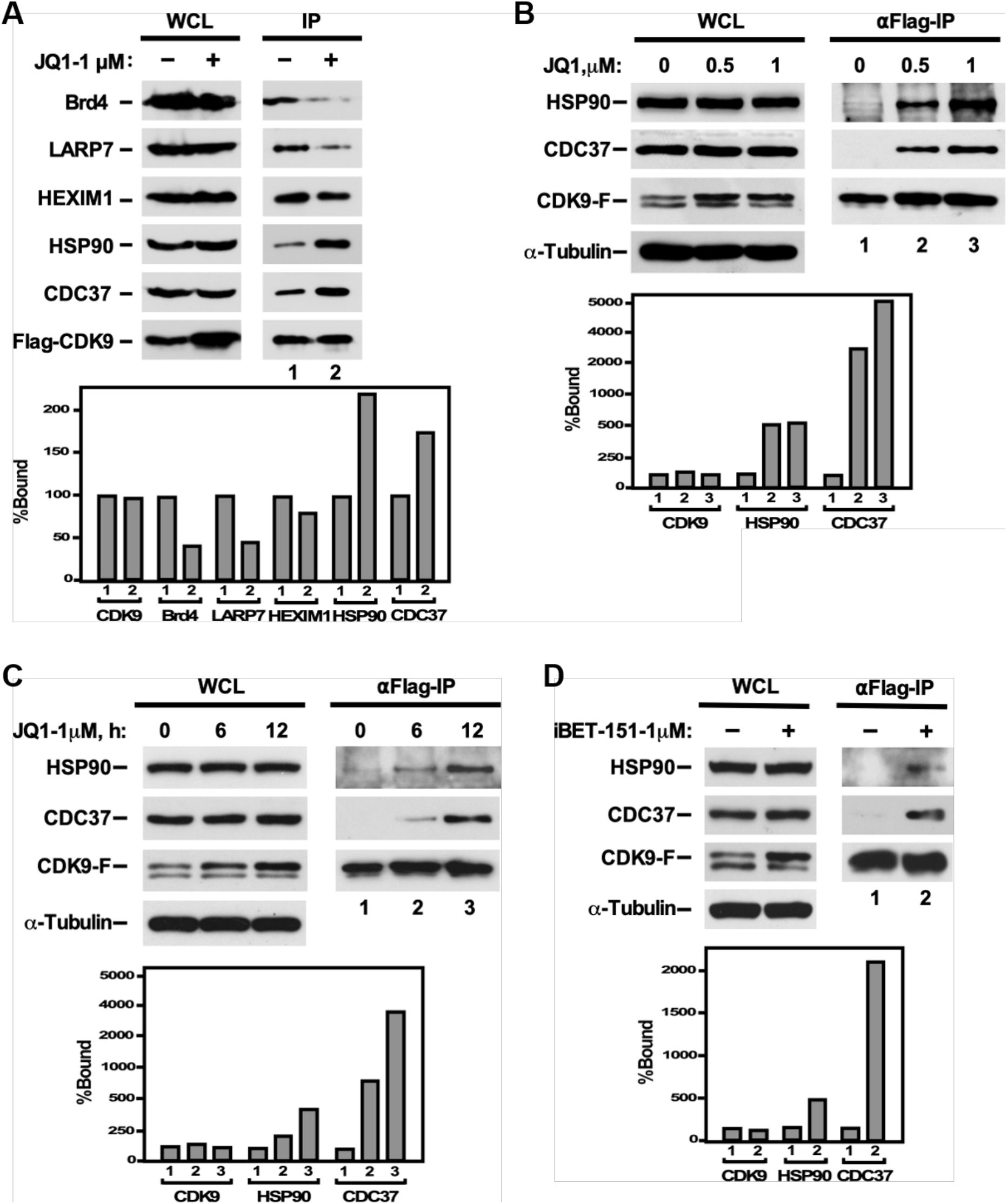
The enhanced association of CDK9 with the HSP90-CDC37 complex under BET inhibitors treatment. **A**. F1C2 cells were treated with 1 μM JQ1 for 24 h. Whole cell lysates (WCL) were subjected to anti-Flag immunoprecipitation and then analyzed by western blotting (WB) for the indicated proteins on the left. The indicated proteins bound to the immunoprecipitated CDK9-Flag were quantified, normalized and displayed in the bottom panel of A, with the signals of lane 1 artificially set to 100%. **B**. F1C2 cells were treated with 0 μM, 0.5 μM or 1 μM JQ1 for 12 h. **C**. F1C2 cells were treated with 1 μM JQ1 for 0 h, 6 h or 12 h. **D**. F1C2 cells were treated with 1 μM iBET-151 for 12 h. The WB results of B, C and D were analyzed and quantified as A.

### The HSP90-CDC37-P-TEFb complex is required for BETi-induced HIV-1 transactivation

To evaluated the functional significance of the HSP90-CDC37-P-TEFb complex for BETi-induced HIV-1 transactivation, we knocked down HSP90 or CDC37 in a HeLa-based NH1 cell line model containing the integrated HIV-1 long terminal repeat (LTR)-driven luciferase reporter gene.^22^ The analysis by WB and qRT-PCR revealed an up to 70% decrease in cellular HSP90 or CDC37 level by the specific shRNA (Figure 3A, B, C&D). In the HSP90 knockdown (KD) NH1 cells, the JQ1-induced Tat-dependent or independent HIV-1 LTR activity was reduced by approximately 60% (Figure 3E). The same phenomenon was also observed in the CDC37 KD cells (Figure 3F). These results indicate that the formation of the HSP90-CDC37-P-TEFb complex could affect the JQ1-induced HIV-1 transcription, suggesting that the complex plays a key role in regulating HIV-1 LTR function once Brd4’s activity is blocked.

**Figure 3.**
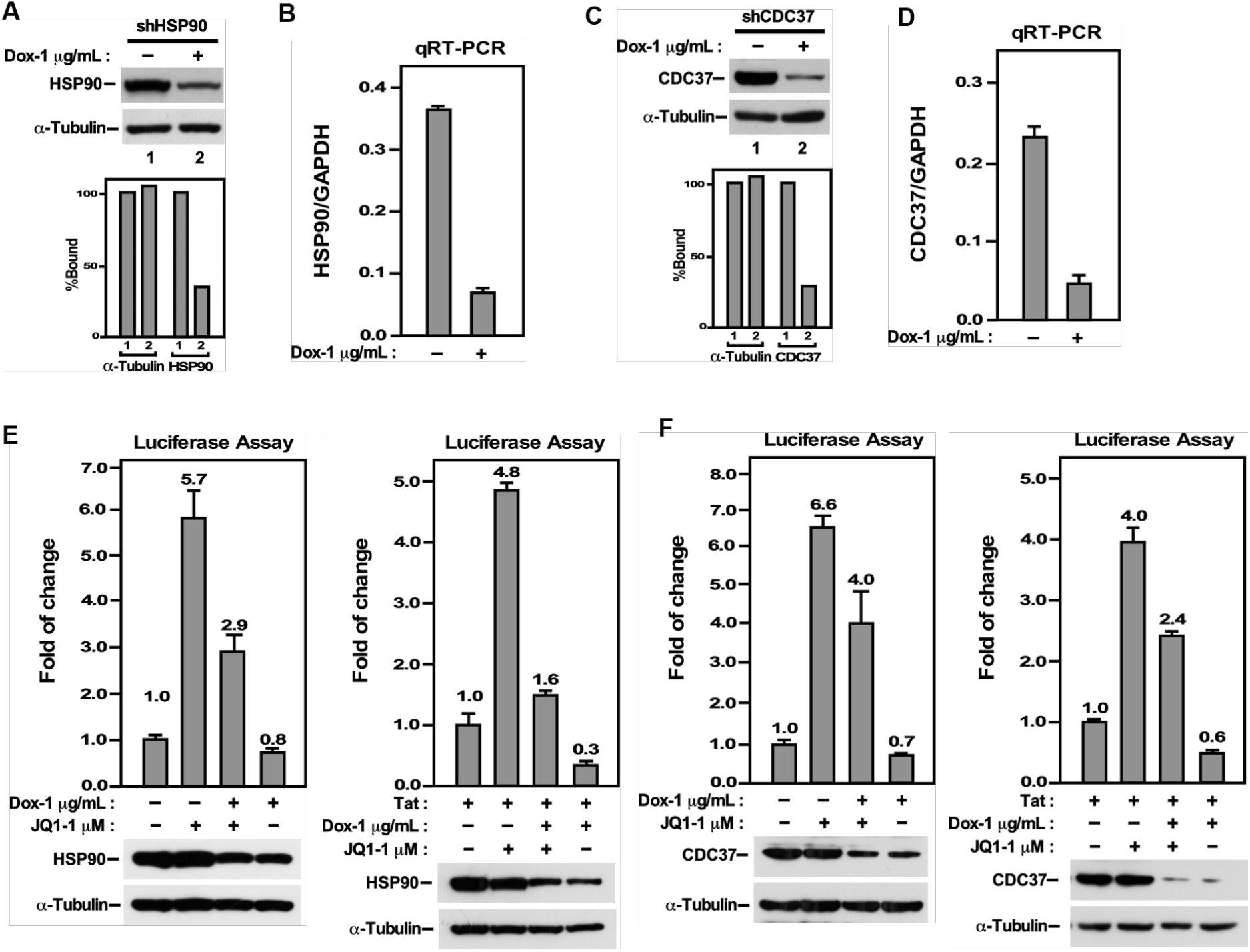
The HSP90-CDC37-CDK9 complex is required for JQ1-induced HIV-1 reactivation. **A, B, C&D**. NH1 cells were transfected with lentivirus harboring either HSP90-targeted or CDC37-targeted shRNA. The efficiency of knocking down HSP90 or CDC37 was assessed by Western blotting (WB) or qRT-PCR of cells induced by 1 μg/mL doxycycline for 72 h. The results of WB were shown in A or D. The binding of CDC37 or HSP90 in lane 2 were quantified with normalization to the signals of tubulin and displayed in the bottom panel of A or D. The signals of lane 1 was set to 100%. **B&D**. The mRNA levels of CDC37 or HSP90 were analyzed by RT-qPCR and normalized to those of GAPDH. The error bars represent mean +/- SD from three independent measurements. **E&F**. The NH1 cells expressing shHSP90 or shCDC37 were induced with 1 μg/mL doxycycline for 60 h, followed by treating cells with 1 μM JQ1 for 12 h. WCL were examined for luciferase activities (top panel) and WB for the indicated proteins (bottom panel).

### Formation of the HSP90-CDC37-P-TEFb complex depends on HSP90’s ATPase activity

We have previously reported that CDK9 is a client protein of HSP90-CDC37, which is essential for P-TEFb’s maturation and stability.^18^ HSP90 normally relies on its ATPase activity to regulate the folding of its client protein.^25^ Notably, when the HSP90 ATPase activity was inhibited by 17-AAG,^26^ the JQ1-induced interaction between HSP90-CDC37 and P-TEFb was significantly disrupted (Figure 4A). Similarly, the JQ1-induced HIV-1 transcription in NH1 and NH2 cells was also inhibited by 17-AAG (Figure 4B&C). The same results were observed with another HSP90 inhibitor, Radicicol (Figure 4D, Figure 4E&F).^27^ These results show that the ATPase activity of HSP90 was necessary for the HSP90-CDC37-P-TEFb complex formation, and that the disruption of this complex caused significant inhibition of the JQ1-induced HIV-1 transactivation. Thus, formation of the HSP90-CDC37-P-TEFb complex has a positive effect on the ability of BETi to induce HIV-1 transactivation.

**Figure 4.**
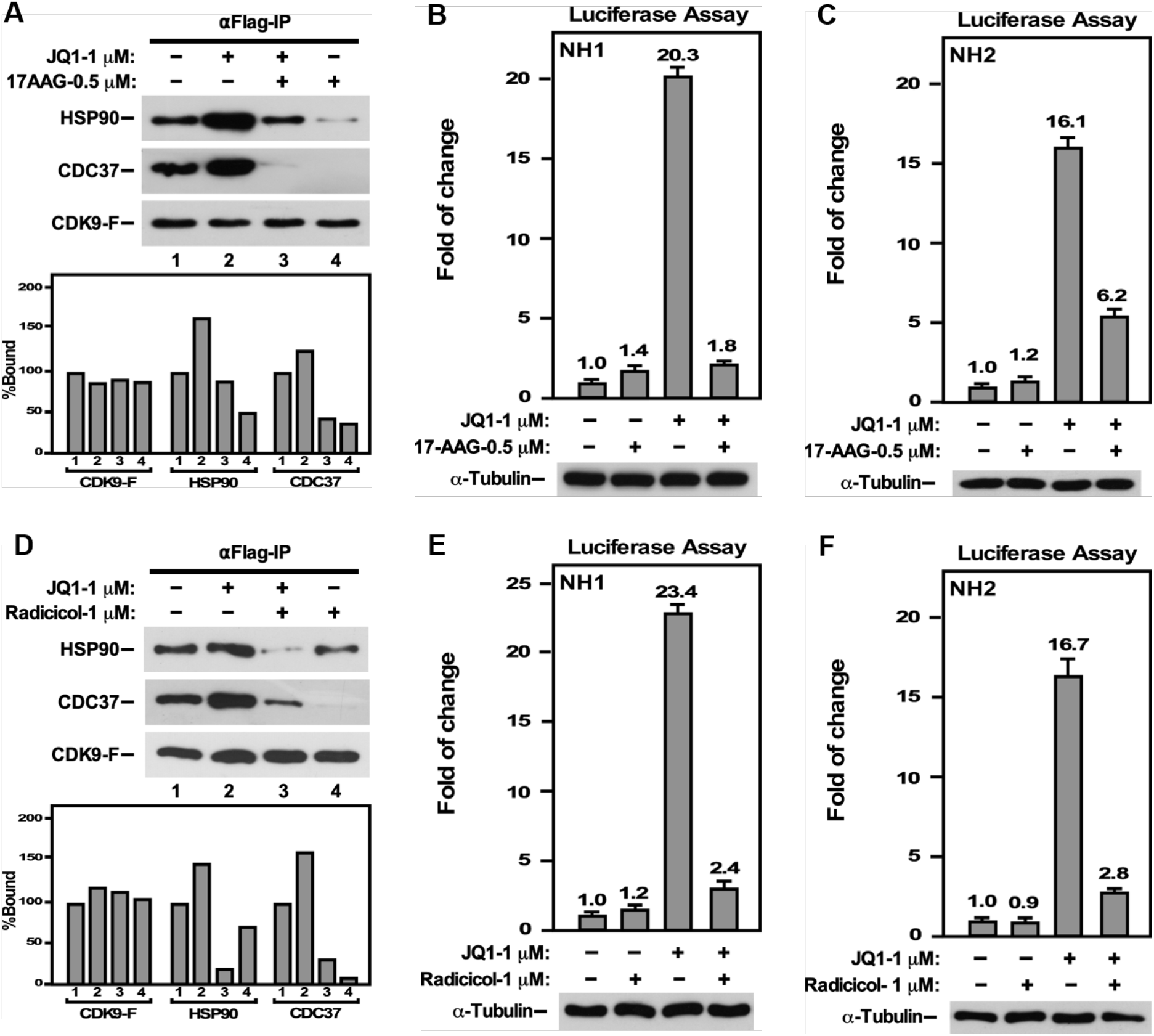
ATPase activity of HSP90 is required for formation of the HSP90-CDC37-P-TEFb complex. **A&D**. F1C2 cells were pre-treated with 0.5 μM 17-AAG or 1 μM Radicicol for 2 h, and then co-treated with 1 μM JQ1 for 12 h. Whole cell lysates (WCL) were subjected to anti-Flag immunoprecipitation (IP), then analyzed by Western blotting (WB) for the indicated proteins on the left. The indicated proteins bound to the immunoprecipitated CDK9-Flag were quantified, normalized and displayed in the bottom panel of A&D. The signals of lane 1 artificially set to 100%. **B, C, E&F**. F1C2 cells were treated in the same way as described in A&C, then analyzed for luciferase activities (top panel) and WB (bottom panel).

### HSP90-CDC37-P-TEFb is recruited by HSF1 to HIV-1 LTR for JQ1 induction of HIV-1 transcription and latency reversal

HSP90-CDC37-P-TEFb has been reported to be recruited by heat shock factor protein 1 (HSF1) to facilitate the proteasome inhibitor-induced HIV-1 reactivation through HSF1’s binding to heat shock elements (HSE) located in the HIV-1 LTR.^28^ In sight of this observation, we examined whether HSF1 participated in JQ1’s induction of HIV-1 transcription. First, an anti-HSP90 IP was performed and the result revealed increased interactions of HSP90 with HSF1 and CDK9 upon JQ1 treatment (Figure 5A). Meanwhile, an anti-HSF1 IP also showed increased interactions of HSF1 with HSP90 and CDK9 (Figure 5B).

**Figure 5.**
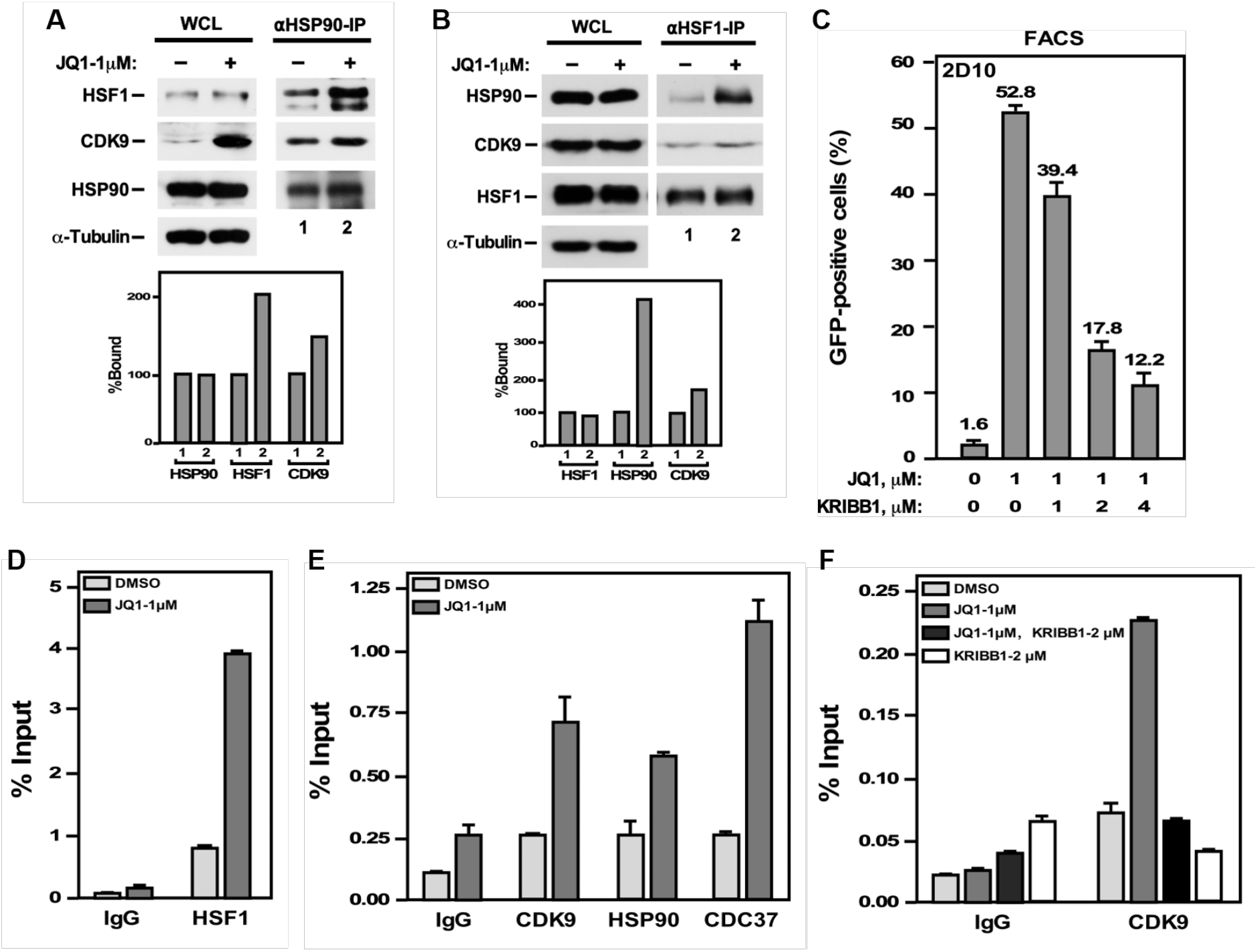
HSF1 recruits more HSP90-CDC37-P-TEFb complex to HIV-LTR promoter for promoting HIV-1 transcription under JQ1 treatment. **A&B**. 2D10 cells were treated with 1 μM JQ1 for 12 h. WCL were subjected to anti-HSP90 or anti-HSF1 immunoprecipitation, and then analyzed by Western blotting for indicated proteins. The indicated proteins bound to the immunoprecipitated HSP90 or HSF1 were quantified, normalized and displayed in the bottom panel of B. The signals of lane 1 were artificially set to 100%. **C**. 2D10 cells were co-treated with 1 μM, 2 μM and 4 μM of HSF1 inhibitor (KRIBB11) and 1 μM JQ1 for 12 h. The induction of GFP expression were analyzed by flow cytometry (FACS). **D, E&F**. 2D10 cells were treated with indicated compounds for 12 h. Whole cell lysates (WCL) were subjected to ChIP analysis to assess the occupancy of HSF1 at HIV-LTR. The signals were normalized to the input DNA. All error bars represent mean +/- SD from three independent measurements.

In addition, the JQ1-induced HIV-1 latency reactivation in Jurkat 2D10 cells, a well-described latency model,^29^ could be inhibited by the HSF1 inhibitor KRIBB11 in a dose-dependent manner (Figure 5C).^30^ Further, the ChIP-qPCR analysis revealed an increase in HSF1’s occupancy at HIV-1 LTR promoter in JQ1-treated cells (Figure 5D). In addition, the levels of CDK9, HSP90 and CDC37 were also increased at the HIV-1 LTR upon JQ1 treatment (Figure 5E). However, when cells were pretreated with the HSF1 inhibitor, the JQ1-induced increase of CDK9’s occupancy was blocked (Figure 5F). Based on these observations, we hypothesized that the inhibition of Brd4 function by JQ1 causes more HSF1 to bind to the HIV-1 LTR promoter, leading to the recruitment of more HSP90-CDC37-P-TEFb complex to promote viral transcription.

### HSP90-CDC37-P-TEFb complex is involved in the equilibrium of P-TEFb in cells

Up to 90% of cellular P-TEFb are reported to be sequestrated in the inactive 7SK snRNP, which is known as a critical factor for maintaining HIV-1 latency.^31^ Like many latency-reversing agents (LRAs), JQ1 may also cause dissociation of 7SK snRNP to release more active CDK9 to promote latent HIV-1 transcription (Figure 1C).^11^ Thus, we speculated that the JQ1-induced formation of the HSP90-CDC37-P-TEFb complex could participate in the dynamic redistribution of active CDK9 in a cell. Indeed, as shown in Figure S2, JQ1 induced disassociation of 7SK snRNP in F1C2 cells. However, more LARP7 and HEXIM1 became associated with CDK9-Flag when HSP90 or CDC37 was knocked down in F1C2 cells (Figure S2A&B). These observations suggest that the disruption of the HSP90-CDC37-P-TEFb complex may promote the reformation of 7SK snRNP. Furthermore, the induced disassociation of 7SK snRNP by JQ1 was reversed when cells were pretreated with HSP90 inhibitors (Figure S2C&D), indicating that the HSP90-CDC37-P-TEFb complex is an important component of the P-TEFb regulatory network in cells.

## Discussion

The chromatin adaptor protein Brd4 binds to acetylated histones and key components of the transcriptional machinery to participate in a number of steps of gene transcription, include chromatin decompaction,^2, 32-34^ formation of the transcriptional initiation complex and transcription elongation.^4, 5^ BETi that act on Brd4 have been studied extensively for their potential effects in treating various kinds of cancers.^35-38^ It is interesting to note that the first BETi, JQ1, has also been found to effectively activate latent HIV transcription via antagonizing Brd4 to promote the formation of the Tat-SEC complex without inducing global T cell activation.^12^ Although JQ1 was shown to induce the release of P-TEFb from 7SK snRNP, which weakly activates basal HIV transcription in Jurkat cells, the mechanism has not been systematically studied.

Through performing quantitative proteomic analysis of the CDK9 interactome in JQ1-treated cells, we found that CDK9 could be recruited to form the HSP90-CDC37-P-TEFb complex. Furthermore, the BETi-induced HIV-1 reactivation could be suppressed by knocking down HSP90 or CDC37. The HSP90-CDC37-P-TEFb complex-mediated HIV-1 transcription could be inhibited by two HSP90 inhibitors 17-AAG and Radicicol that suppress the ATPase activity of HSP90. The results confirmed the formation of the HSP90-CDC37-P-TEFb complex, which is closely involved in HIV-1 latency reactivation. Moreover, HSF1 has been identified as playing a mediating role to recruit the HSP90-CDC37-P-TEFb complex to the HIV-1 promoter for activated transcription. Thus, the HSF1-HSP90-CDC37-P-TEFb axis may compensate for the loss of Brd4-P-TEFb to activate HIV transcription.

In human cells, most P-TEFb are known to be sequestrated in the 7SK snRNP in an inactive state.^39^ The active P-TEFb that exist outside of this snRNP can be competitively recruited to the HIV-1 promoter by Brd4 (Brd4-P-TEFb complex),^4, 40^ or by HIV Tat (Tat-SEC complex).^41^ Our study revealed that there exists an equilibrium between several CDK9-containing complexes including the Brd4-P-TEFb complex, 7SK snRNP, the SEC complex and then the HSP90-CDC37-P-TEFb complex once the Brd4 function is eliminated (Figure 6).

**Figure 6.**
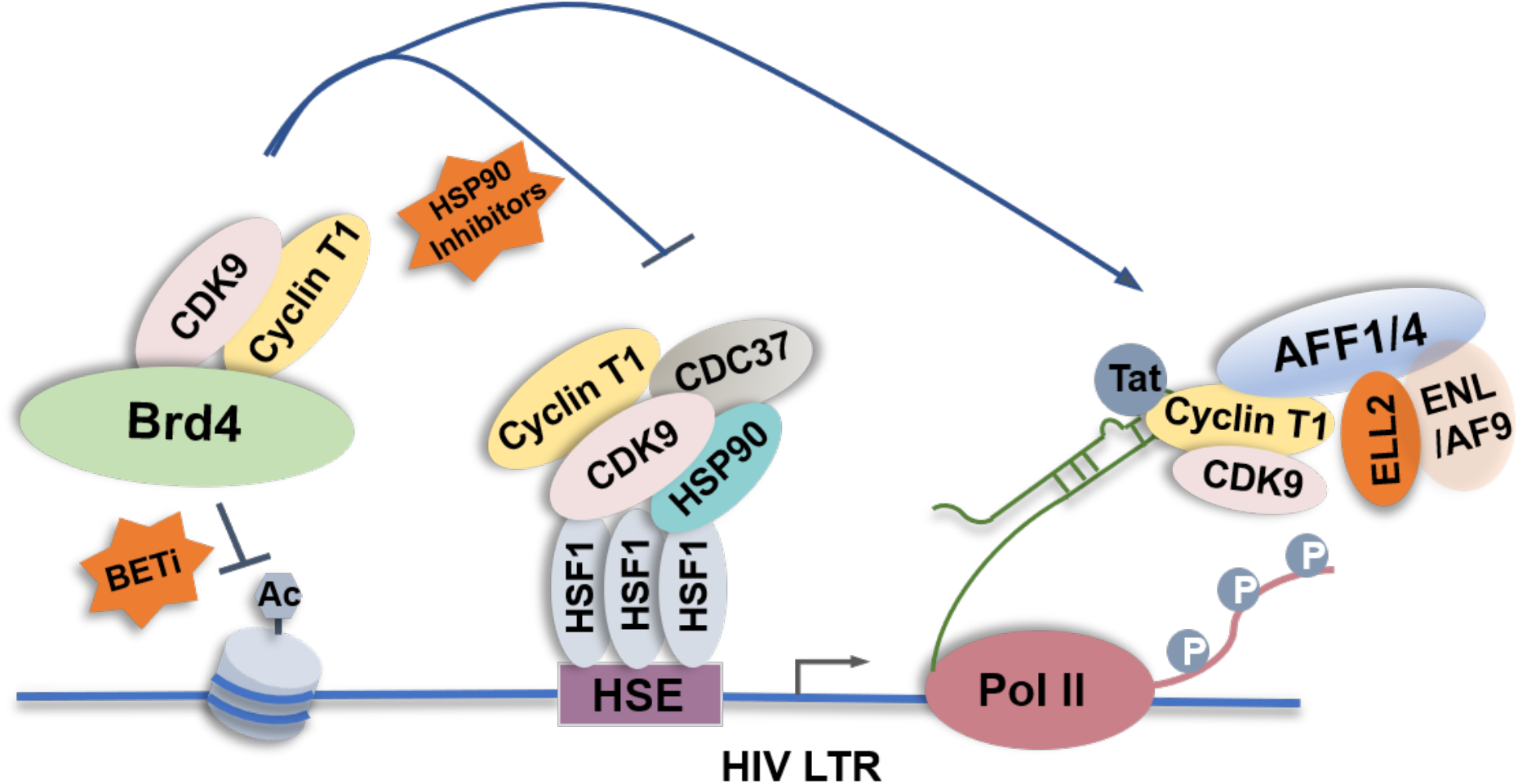
Schematic representation of the HSP90-CDC37-P-TEFb pathway activated by Brd4 inhibition for HIV-1 reactivation.

Recently, two groups have reported different latency-reversing agents (LRAs) to activate latent HIV in animal models,^42, 43^ which may enable the strategy nick-named “shock and kill” to eliminate latent HIV-1 reservoirs. As a potential LRA, BETi can effectively perturb the latent HIV reservoir in cell models. Beside the activation of Tat-dependent HIV transcription, BETi JQ1 induces the HSP90-CDC37-P-TEFb complex formation to promote HIV transcription by a HSF1-dependent mechanism. This compensatory mechanism between HSP90-CDC37-P-TEFb and Brd4-P-TEFb demonstrates the importance of HSP90 in regulating HIV transcription. It is worth noting that several HSP90 inhibitors are currently in clinical trials for treating solid malignancies and lymphomas.^44^ These inhibitors can prevent T-cell activation, cytokine production and drug resistance, which have been considered to be of great promise for treatment of latent HIV infection.^45^ Thus, the novel mechanism reported here suggests the rationality of the concomitant use of multiple drug targets for achieving viral eradication in the future.

## Conclusion

The mass spectrometry-based quantitative proteomics provides a comprehensive tool to systematically study the dynamic association and disassociation of intrinsic protein complexes that control transcriptional elongation under diverse conditions. Our quantitative analysis of the CDK9 interactome under BETi treatment conditions demonstrates that the interaction of the HSP90-CDC37 complex with CDK9 was significantly increased for regulation of HIV-1 gene transcription in BETi-treated cells. Formation of the HSP90-CDC37-P-TEFb complex could not only stabilize the protein level of CDK9, but also compensate for the loss of Brd4-P-TEFb to activate HIV transcription through the interaction with HSF1 that recruits the complex to the viral LTR. Several key CDK9-containing complexes, including Brd4-P-TEFb, 7SK snRNP, and HSP90-CDC37-P-TEFb, could synergistically regulate the function of P-TEFb in an equilibrium that is key for regulating transcriptional elongation. Our results suggest a new strategy for eradicating latent HIV through using combinations of latency-reversing agents and the anti-viral inhibitors.

## Associated Content

### Supporting Information

The supporting Information is available free of charge on the ACS Publications website at Figure S1. CDK9 interactome analysis; Figure S2. JQ1 partially disassociates CDK9 form 7SK snRNP to form HSP90-CDC37-P-TEFb complex; Extended Data 1. The list of proteins identified from replicate 1 and 2; Extended Data 2. The list of annotated sequence for identification and quantification of CDK9, HSP90 alpha, HSP90 beta and CDC37.

## Author Contributions

X.G., Y.X., and Q.Z. conceived and designed the study. C.W., Z.P., and Y.H. conducted sample acquisition and extracted the protein, performed protein digestion, SILAC labeling, HPLC-MS/MS analysis, and FCAS analysis. C.W., Z.P., Z.Z., and R.L. performed the IP and WB experiments for data validation. C.W., X.G., and Q.Z. prepared the manuscript. All authors have given approval to the final version of the manuscript.

## Notes

The authors declare no conflict of interest.

## Acknowledgement

This work was supported by the National Key Research and Development Program of China (grant 2020YFA0608300), the National Natural Science Foundation of China (grant 81672955), the Xiamen Southern Oceanographic Center (grant 17GYY002NF02), and the National Institutes of Health (grant R01AI41757).

## Materials and methods

### Reagents and cell cultures

Inhibitors used in this study were Radicicol (Abcam, Cat# ab141848), 17-AAG (MedChemExpress, Cat# HY-10211), KRIBB11 (MedChemExpress, Cat# HY-100872), JQ1 (Cayman, Cat# 11187), and iBET-151 (MedChemExpress, Cat# HY-13235). Antibodies used in this study were HSP90 (Abcam, Cat# ab133491), CDC37 (Abcam, Cat# ab109419), HSF1 (Cell Signaling Technology, Cat# 4356), Cyclin T1 (Santa Cruz Biotechnology, Cat# sc-10750); Anti-BRD4, AFF4, LARP7, HEXIM1 and CDK9 were generated in our own laboratory as described previously.^4, 46^ HeLa cells, NH1 cells and F1C2 cells were cultured with Dulbecco’s Modified Eagle Medium (DMEM) with 10% fetal bovine serum (FBS). 2D10 cells were cultured in RPMI-1640 supplemented with 10% FBS.

### Flow cytometry analysis

The flow cytometry-based experiments were performed in 2D10 cells treated with the indicated concentrations of HSP90 inhibitors (17-AAG or Radicicol), HSF1 inhibitor KRIBB11, and JQ1 for the indicated time. After harvesting, 2D10 cells were washed twice with 1 x phosphate buffered saline (PBS) and analyzed by flow cytometer (Epics Altra, BECKMAN COULTER) for the percentages of cells expressing GFP.

### Luciferase reporter assay

The HeLa-based NH1 cells were used for luciferase assay containing HIV-LTR driven luciferase reporter gene. NH1 cells were treated with indicated concentrations of different inhibitors for 12 h or for indicated time. Whole cell lysates were prepared and luciferase activity was measured using a kit from Promega.

### Quantitative RT-PCR assay

Total RNA from NH1 cells were extracted using Trizol agent and reverse transcribed with random primers. The cDNAs were subjected to quantitative real time PCR with the following primers:

HSP90AA1:

F: 5’-AACGATGATGAGCAGTACGCTTG-3’

R: 5’-TGTTCCACGACCCATAGGTTCA-3’

HSP90AB1:

F: 5’-GAGAACCTCTGCAAGCTCATGAAAG-3’

R: 5’-GCAGCAAGGTGAAGACACAAGTCTA-3’

### shRNA knockdown (KD) of HSP90 in NH1 cells

The NH1 cells were infected with the pLKO.1-puro-based lentiviral vector containing following sequences:

shHSP90AA1:

F: TATGGCATGACAACTACTTTACTCGAGTAAAGTAGTTGTCATGCCATA-3’.

R: ATGGCATGACAACTACTTTACTCGAGTAAA GTAGTTGTCATGCCATA-3’.

shHSP9AB1:

F: 5’-CGCATGGAAGAAGTCGATTAGCTCGAG TAATCGACTTCTTCCATGCG-3’.

R: 5’-CGCATGGAAGAAGTCGATTAGCTCGAGCTAATCG ACTTCTTCCATGCG-3’.

Two days after infection, the cells were induced by 1 μg/mL doxycycline for 72 hours. The KD efficiencies of the pools were examined by WB.

### Co-immunoprecipitation (co-IP) assays

Whole cell lysates or nuclear extracts were prepared from F1C2 cells or 2D10 cells. Anti-Flag immuno-precipitate was incubated with anti-Flag agarose beads for 3 h or overnight. Anti-HSF1 immuno-precipitate was incubated with anti-HSF1 antibody overnight and then incubated with protein-A/G beads for 2 h. After incubation, anti-Flag agarose or protein-A/G beads were washed extensively with buffer D0.15 (15% Glycerol, 0.2% NP40, 20 mM HEPES pH 7.9, 0.2 mM EDTA-2Na, 0.15 M KCl, 1 mM DTT, 1 mM PMSF) and buffer D0.1 (15% Glycerol, 0.2% NP40, 20 mM HEPES pH 7.9, 0.2 mM EDTA-2Na, 0.1 M KCl, 1 mM DTT, 1 mM PMSF). In the case of anti-Flag agarose beads, proteins were competitively eluted with Flag peptides, while for protein-A/G beads, proteins were eluted with glycine at pH 2.0. Finally, eluted proteins were analyzed by WB using indicated antibodies.

### Chromatin immunoprecipitation (ChIP) assay

After treating with JQ1 for 12 h, 2D10 cells were cross-linked with 1% formaldehyde for 5 min at room temperature. Post-washing with PBS, 1 × 10^7^ cells were re-suspended with 300 μL SDS lysis buffer (1% SDS, 10 mM EDTA, 50 mM Tris-HCl, pH 8.0), followed by sonication (30 sec ON, 10 sec OFF, for a total of 35 cycles). The lysates were diluted 10 times with dilution buffer (0.01% SDS, 1.1% Triton X-100, 1.2 mM EDTA, 16.7 mM Tris-HCl, pH 8.1, and 167 mM NaCl). Incubated with antibodies overnight at 4°C. After mixing with Protein A/G beads for 1 h, the beads were washed thrice with the wash buffer (0.1% SDS, 1% Triton X-100, 2 mM EDTA, 20 mM Tris-HCl, pH 8.1, and 150 mM NaCl) and then washed twice with the TE buffer (10 mM Tris-HCl, pH 8.1 and 1 mM EDTA). DNA was eluted from the beads using the elution buffer (1% SDS and 100 mM NaHCO_3_) and purified by PCR purification kit (Promega). The final products were analyzed by quantitative real-time PCR with the primers listed below:

HIV-1 LTR:

F: 5’-GTTAGACCAGATCTGAGC CT-3’.

R: 5’-GTGGGTTCCCTAGTTAGCCA-3’.

The PCR signals were normalized to the input DNA and shown as an average of three independent measurements.

### Quantitative proteomics analysis of CDK9 interactome

F1C2 cells were cultured by SILAC DMEM supplemented with ^13^C_6_-Arginine [Arg^6^], ^13^C_6_-Lysine [Lys^6^], and 10% dialyzed FBS for 7 times of passage. The heavy cells were treated with JQ1 for 12 h. After mixing with same amount of normal F1C2 cells treated with DMSO, the cell mixtures were lysed with buffer containing 50 mM Tris-HCl (pH 7.9), 150 mM NaCl, 1 mM EDTA and 1% NP-40. Then, cell lysates were pooled and subjected to affinity purification by using anti-Flag M2-agarose beads. The beads were washed extensively with buffer containing 50 mM Tris-HCl (pH 7.9), 150 mM NaCl, 1 mM EDTA and 0.2% NP-40, and eluted with 1 × Flag peptides (Sigma).

Eluted proteins were subjected to in-solution digestion according to FASP method.^47^ Briefly, the eluted proteins of IP were transferred to a 10 kD centrifugal spin filter (Millipore), and followed by washing with 8 M urea 200 μL for three times. Subsequently the samples were reduced with 50 mM dithiothreitol (DTT) at room temperature (RT) for 30 min, alkylated with 50 mM iodoacetamide (IAA) at RT for 30 min in the dark, and followed by washing with 8 M urea 200 μL for three times by centrifugation at 20 °C with 13000 *g*. Then, trypsin dissolved in 50 mM tetraethylammonium carbonate (TEAB) was added to the filter to digest samples at 37 °C for 16-18 hours (trypsin/substrate, m/m = 1:50). The digested samples were transferred to a new collection tube by centrifugation and desalted by StageTips filled with C18 beads. For sample desalting, StageTips were firstly equilibrated with 200 μL acetonitrile (ACN) for three times and 200 μL 0.1% formic acid (FA) for three times. Then, the tips were applied digested samples and washed with 200 μL 0.1% FA for three times. After eluting with 0.1% FA/75% ACN, samples were concentrated with vacuum centrifugation and store at -20 °C for further analysis.

The tryptic peptides were resuspended in 0.1% formic acid (FA)/2% acetonitrile (ACN) and subjected to a nanoscale UHPLC system connected to an Orbitrap Q-Exactive mass spectrometer (Thermo Fisher Scientific). The peptides were separated on a 75 μm × 15 cm RP-HPLC analytical column packed with 2 μM C18 particles (Thermo Fisher Scientific) using an increasing gradient of 0.1% FA in ACN (8-35% over 100 min) at a flow rate of 300 nL/min. The mass spectrometer was operated in a profile mode (resolution: 70,000; automatic gain control (AGC) value: 3e6; maximum injection time: 50 ms) at full MS scan over a mass range between *m/z* 350 and 1800. The twenty most intense ions in each MS survey scan were automatically selected for MS/MS (resolution: 17500; AGC value: 1e5; maximum injection time: 60 ms). The dynamic exclusion window was set at 50 s. Raw LC-MS/MS data were analyzed with Proteome Discoverer software 2.4 (Thermo Fisher Scientific). The spectrum selector was set to its default values to search data against the uniport_sprot human proteome database. The precursor mass tolerance was set to 10 ppm. Fragment mass tolerance was set to 20 mmu. Oxidation (Met) (+15.9949 Da) and acetylation (protein N-terminus) (+42.0106 Da) were set as variable modifications. For protein identification, two missed cleavages were allowed and at least one unique peptide was required. Ontology enrichment analysis was performed using Metascape^48^.

## Notes

### Competing Interest Statement

The authors have declared no competing interest.

