## Supporting information for "Quantitative proteomics of the CDK9 interactome reveals a novel function of the HSP90-CDC37-P-TEFb complex for BETi-induced HIV-1 latency reactivation"

**Figure S1. CDK9 interactome analysis. A.** Correlation analysis of two biological replicates. Scatter plot illustrating the log2 ratio of replicate1 and replicate2. **B.** Enriched ontology clusters of CDK9-interacted proteins (n= 535)**. C, D, E&F.** MS/MS spectra of the representative peptides of CDK9 (C), HSP90 alpha (D), HSP90 beta (E) and CDC37 (F).

**Figure S2. JQ1 partially disassociates CDK9 form 7SK snRNP to form HSP90-CDC37-P-TEFb complex. A.** shHSP90 or shCDC37 were overexpressed in F1C2 cells by inducing with 1 μg/mL doxycycline for 72 h. Whole cell lysates (WCL) were subjected to anti-Flag immunoprecipitation (IP), then analyzed by WB for the indicated proteins on left. The indicated proteins bound to the immunoprecipitated CDK9-Flag in lanes 2, 3 and 4 of B were quantified, normalized to the signals of CDK9-Flag and displayed in the bottom panel of A&B. The signals of lane 1 artificially was set to 100%. **C&D.** F1C2 cells were pre-treated with 0.5 μM HSP90 inhibitor 17-AAG or 1 μM Radicicol for 2 h and then treated with 1 μM JQ1 for 12 h. Anti-Flag immunoprecipitation derived from WCL was measured by Western blotting for the indicated proteins. The bound of these proteins were quantified as A (bottom panel).

**Extended Data 1. The list of proteins identified from replicate 1 and 2.**

**Extended Data 2. The list of annotated sequence for identification and quantification of CDK9, HSP90 alpha, HSP90 beta and CDC37.**

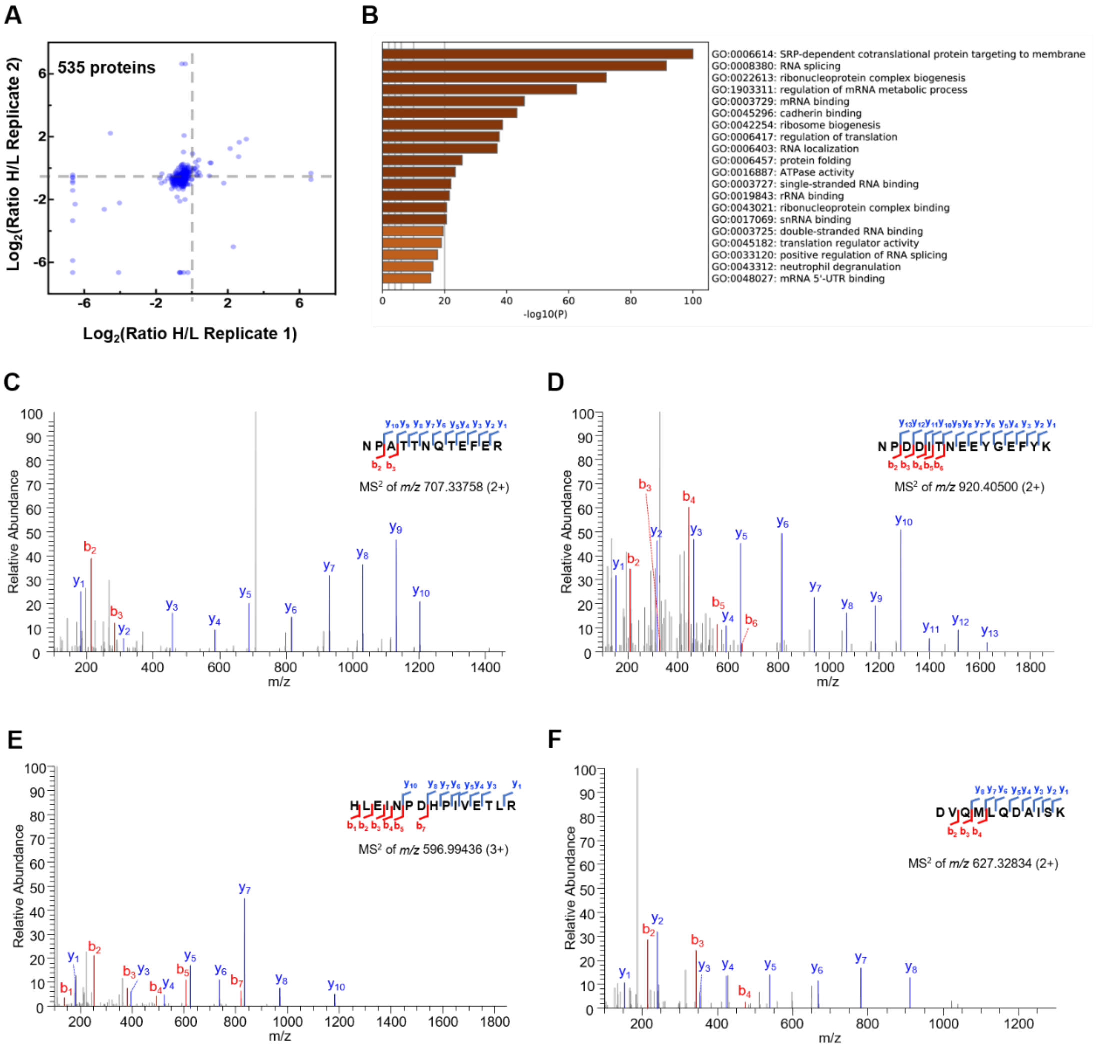
A
535 proteins
Log2(Ratio H/L Replicate 2)
Log2(Ratio H/L Replicate 1)
B
GO:0006614: SRP-dependent cotranslational protein targeting to membrane
GO:0008380: RNA splicing
GO:0022613: ribonucleoprotein complex biogenesis
GO:1903311: regulation of mRNA metabolic process
GO:0003729: mRNA binding
GO:0045296: cadherin binding
GO:0042254: ribosome biogenesis
GO:0006417: regulation of translation
GO:0006403: RNA localization
GO:0006457: protein folding
GO:0016887: ATPase activity
GO:0003727: single-stranded RNA binding
GO:0019843: rRNA binding
GO:0043021: ribonucleoprotein complex binding
GO:0017069: snRNA binding
GO:0003725: double-stranded RNA binding
GO:0045182: translation regulator activity
GO:0033120: positive regulation of RNA splicing
GO:0043312: neutrophil degranulation
GO:0048027: mRNA 5'-UTR binding
-log10(P)
C
NPATTNQTEFER
MS2 of m/z 707.33758 (2+)
Relative Abundance
m/z
D
NPDDITNEEYGEFYK
MS2 of m/z 920.40500 (2+)
Relative Abundance
m/z
E
HLEINPDHPIVETLR
MS2 of m/z 596.99436 (3+)
Relative Abundance
m/z
F
DVQMLQDAISK
MS2 of m/z 627.32834 (2+)
Relative Abundance
m/z

**Figure S1**

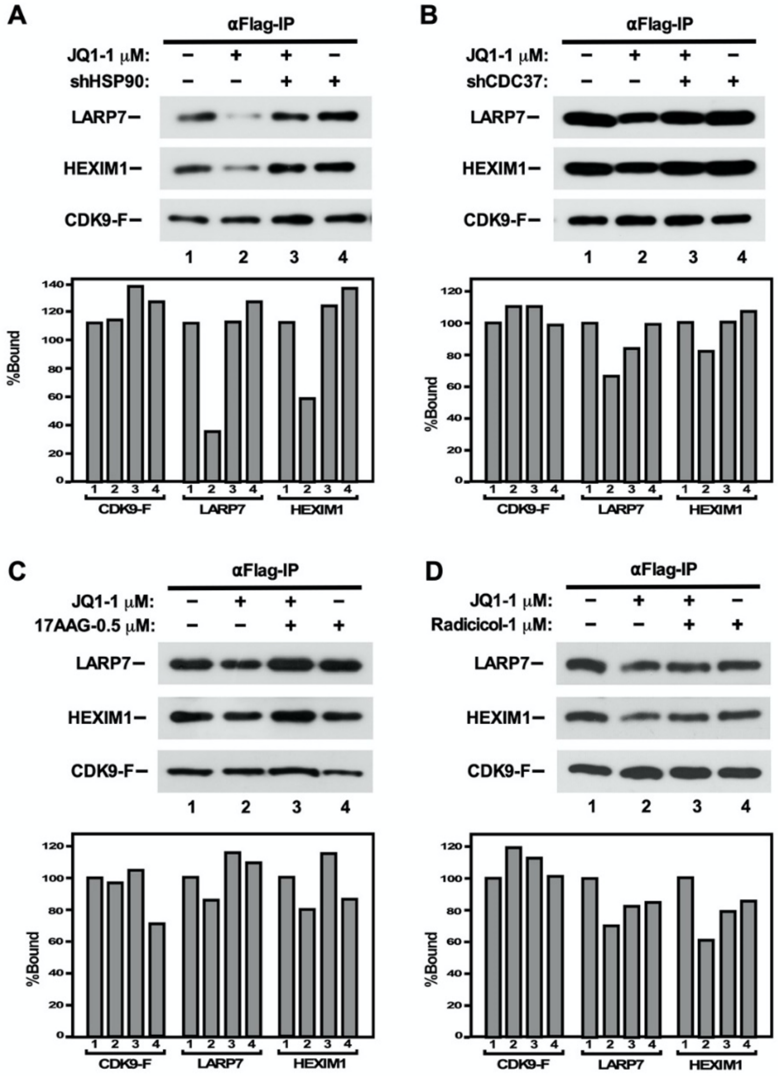
A
αFlag-IP
JQ1-1 μM: – + + –
shHSP90: – – + +
LARP7
HEXIM1
CDK9-F
1 2 3 4
%Bound
CDK9-F
LARP7
HEXIM1
B
αFlag-IP
JQ1-1 μM: – + + –
shCDC37: – – + +
LARP7
HEXIM1
CDK9-F
1 2 3 4
%Bound
CDK9-F
LARP7
HEXIM1
C
αFlag-IP
JQ1-1 μM: – + + –
17AAG-0.5 μM: – – + +
LARP7
HEXIM1
CDK9-F
1 2 3 4
%Bound
CDK9-F
LARP7
HEXIM1
D
αFlag-IP
JQ1-1 μM: – + + –
Radicicol-1 μM: – – + +
LARP7
HEXIM1
CDK9-F
1 2 3 4
%Bound
CDK9-F
LARP7
HEXIM1

**Figure S2**
